# Neural substrates of top-down processing during perceptual duration-based timing and beat-based timing

**DOI:** 10.1101/2023.06.06.543032

**Authors:** Mitsuki Niida, Yusuke Haruki, Fumihito Imai, Kenji Ogawa

## Abstract

Temporal context is a crucial factor in timing. Previous studies have revealed that the timing of regular stimuli, such as isochronous beats or rhythmic sequences (termed beat-based timing), activated the basal ganglia, whereas the timing of single intervals or irregular stimuli (termed duration-based timing) activated the cerebellum. We conducted a functional magnetic resonance imaging (fMRI) experiment to determine whether top-down processing of perceptual duration-based and beat-based timings affected brain activation patterns. Our participants listened to auditory sequences containing both single intervals and isochronous beats and judged either the duration of the intervals or the tempo of the beats. Whole-brain analysis revealed that both duration judgments and tempo judgments activated similar areas, including the basal ganglia and cerebellum, with no significant difference in the activated regions between the two conditions. In addition, an analysis of the regions of interest revealed no significant differences between the activation levels measured for the two tasks in the basal ganglia as well as the cerebellum. These results suggested that a set of common brain areas were involved in top-down processing of both duration judgments and tempo judgments. Our findings indicate that perceptual duration-based timing and beat-based timing are driven by stimulus regularity irrespective of top-down processing.

## Introduction

Timing is a fundamental ability for humans that occurs in various contexts, and the mechanisms underlying timing can differ depending on the context (Buhusi and Meck 2005; Paton and Buonomano 2018). Among these contexts, the temporal regularity (whether the intervals are regular or irregular) of short intervals is a notable example; the timing of regular intervals is more accurate than the timing of irregular intervals (Drake and Botte 1993; Yee et al. 1994; Jones and Yee 1997). Duration-based timing refers to absolute timing based on the durations of individual intervals, which typically occur in processing single intervals or irregular sequences. In contrast, beat-based timing refers to relative timing that uses internal beats formed based on the periodicity or regularity of external sequences, which typically occur in situations featuring rhythmic external stimuli such as music.

Behavioral studies have examined whether timings based on duration, beat, or both underlie the perception of short intervals. However, the findings of these studies are not consistent with each other. Drake and Botte (1993) proposed that a single mechanism underlies both timings of single intervals and sequences. Keele et al. (1989) and Pashler (2001) suggested that an interval timer, a duration-based mechanism, times short intervals and operates in a beat mode for rhythmic events, whereas McAuley and Jones (2003) questioned this theory. Moreover, Schulze (1978) and Rammsayer and Brandler (2004) showed that the interval timer did not process rhythmic patterns.

A large body of neuroscientific evidence stating that perceptual duration-based timing and beat-based timing have distinct neural substrates has been obtained from several human studies (Breska and Ivry 2016; Paton and Buonomano 2018). The cerebellum is essential for duration-based timing of single subsecond intervals but not for beat-based timing of regular sequences. A neuropsychological study on patients with cerebellar degeneration reported that these patients exhibited deficits in duration-based timing but not in beat-based timing (Grube et al. 2010a). Furthermore, a previous study demonstrated that participants who had transcranial magnetic stimulation applied over their cerebellum exhibited a similar tendency to the patients with cerebellar degeneration (Grube et al. 2010b). On the other hand, normal functioning of the basal ganglia is necessary for beat processing (Grahn 2009). Studies using functional magnetic resonance imaging (fMRI) demonstrated that the putamen was more activated when processing beat sequences than when processing irregular sequences (Grahn and Brett 2007; Grahn and Rowe 2009, 2013). Moreover, patients with Parkinson’s disease, a model of basal ganglia dysfunction, showed impaired discrimination of beat sequences compared with controls, while their discrimination of nonbeat sequences was normal (Grahn and Brett 2009).

Further evidence for the existence of distinct neural systems activated by duration-based timing and beat-based timing was revealed by Teki et al. (2011) using an fMRI experiment. Participants were asked to judge whether the last interval of auditory sequences was longer or shorter than the penultimate interval. The stimuli included either regular or irregular sequences, inducing beat-based or duration-based timing, respectively. The results revealed that an olivocerebellar network was selectively activated by the perceptual timing of irregular sequences and a striato–thalamo–cortical network was selectively activated by the perceptual timing of regular sequences. Interestingly, almost all participants did not notice that there were two types of sequences, although their brains processed the stimuli using two different neural networks, suggesting that the difference in the activated neural networks was caused by the difference in stimulus regularity.

The abovementioned studies used stimuli with distinct contexts to compare duration-based timing with beat-based timing. The findings of these studies suggested that the activation of the olivocerebellar network and striato–thalamo–cortical network was dependent on the stimulus context and that the timing behavior in both contexts involved bottom-up processing. Furthermore, the report that the participants did not notice the difference in stimulus context whereas the different brain networks were activated for the stimuli (Teki et al. 2011) suggests that the olivocerebellar and striato–thalamo–cortical networks are not involved in conscious strategies for processing regular or irregular sequences. These findings were contradictory to those of earlier psychophysical studies in which participants reported using different timing strategies for regular and irregular sequences (Yee et al. 1994; Jones and Yee 1997). Thus, it remains unclear how top-down processing, including attention to temporal regularity and the application of different strategies to timing regular and irregular sequences, is related to the distinct neural systems serving duration-based and beat-based timings.

To address this question, we investigated how top-down processing of duration-based and beat-based timings affected the differential activation of task-associated neural systems. To this end, we used fMRI in participants performing duration-based and beat-based timing tasks and compared brain activation levels measured during the two tasks. To equalize the bottom-up inputs while processing the duration-based and beat-based timings, we presented stimuli that included both single intervals and isochronous beats in both types of timing tasks. Participants were instructed to listen to the auditory sequences and judge either only the duration of the single intervals or only the tempo of isochronous beats. We compared the fMRI images of brain activations during the judgments of intervals versus the judgments of beats to assess top-down processing in these tasks.

## Materials and Methods

### Participants

Twenty healthy volunteers (13 males and 7 females; mean age, 22.35 years; range, 20–27 years) participated in the experiment. The number of participants was determined to detect reliable activations based on previous fMRI experiments (Teki et al. 2011). Their sample size was eighteen, and the required sample size calculated from their maximum and minimum effect sizes ranged from four to nineteen (α, 0.05; power, 0.85). All participants were right-handed as assessed using the Edinburgh Handedness Inventory (Oldfield 1971) modified for Japanese participants (Hatta and Nakatsuka 1975). Written informed consent was obtained from each participant. The experimental protocol received approval from the local ethics committee, and the experiments were conducted in accordance with the 1964 Declaration of Helsinki.

### Stimuli

The stimulus used in our experiment was an auditory sequence consisting of 12 clicks (Fig. 1). The clicks were sine waves (900 Hz or 1000 Hz) of 20 ms duration that faded in over the first 5 ms and faded out over the last 5 ms. The sequence contained two single interval stimuli (2 clicks) and two isochronous beat stimuli (4 clicks) arranged in an alternating order. The first interval and first beat were used as the standard stimuli, and the second interval and second beat were used as the comparison stimuli.

**Fig. 1.**
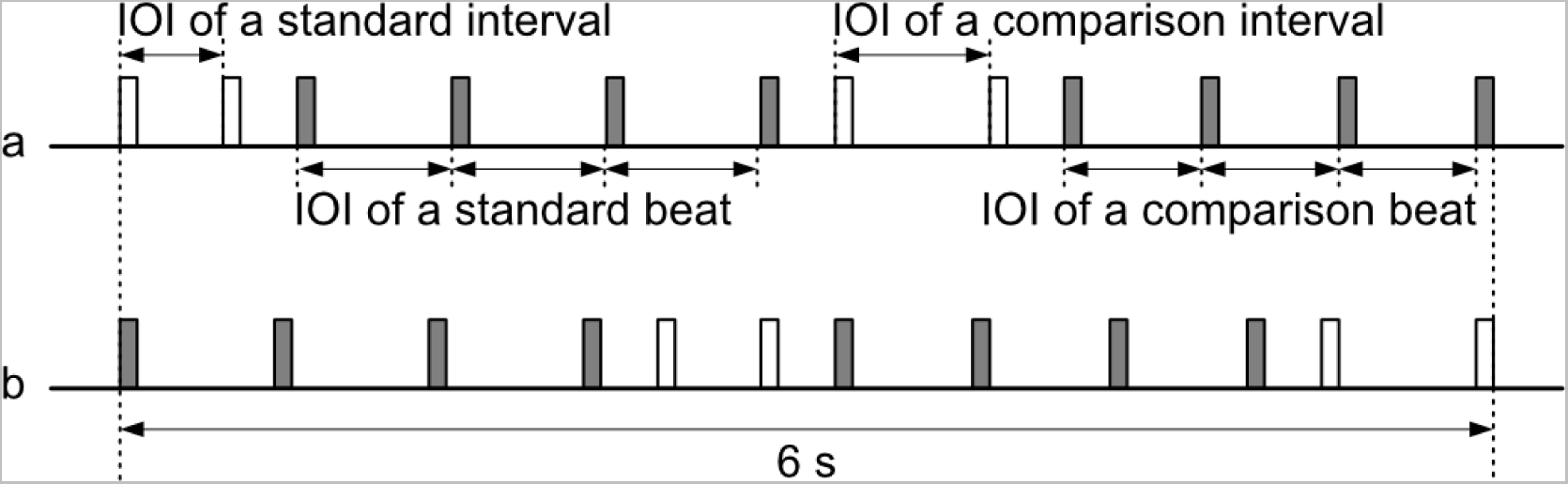
Schematics of the auditory sequences used in the experiments. Each rectangle represents a click sound, with colors representing different sound frequency. The sequence contained two single interval stimuli and two isochronous beat stimuli arranged in an alternating order. The first interval and first beat were used as the standard stimuli, and the second interval and second beat were used as the comparison stimuli. The duration of IOI of the standard interval, standard beat, comparison interval, and comparison beat differed from each other and varied depending on IOI conditions. The total duration of the sequence was 6 s. (a) interval-first sequences. (b) beat-first sequences.

The duration of the inter-onset intervals (IOIs) used for the standard stimuli was 400 ms or 600 ms and differed between the interval and beat stimuli. The ratio of the IOIs used for the comparison stimulus to the standard stimulus varied ±30% for the interval stimulus and ±15% for the beat stimulus. We selected these ratios based on the results of pilot experiments in which IOI ratios were varied to make performance levels comparable for the two stimuli. The conditions and tasks of the pilot experiment were identical to those of the main experiment, except for an IOI ratio condition ranging from 5% to 40% in steps of 5%. These IOI ratios were compared based on the task accuracy, and we selected ±30% for the interval stimulus and ±15% for the beat stimulus, which were the ratios used by Teki et al. (2011), from the ratios that showed no significant difference in the task accuracy.

The click frequency was manipulated to make it easier to distinguish between the intervals and the beats. When the interval stimuli were 900 Hz, the beat stimuli were 1000 Hz. Conversely, when the interval stimuli were 1000 Hz, the beat stimuli were 900 Hz. That is, the intervals and beats in a single trial sequence differed from each other in three ways: (i) IOI of the standard stimulus (400 ms or 600 ms); (ii) IOI ratio of the comparison stimulus (longer, shorter); and (iii) click frequency (900 Hz or 1000 Hz). In addition, (iv) the interval and beat stimuli were presented in a different order (interval first or beat first). Thus, in total, there were 2^4^ = 16 types of sequences, which were presented in random order. The total length of the sequence was 6 s. In each trial sequence, the three intervals between the single interval stimuli and the isochronous beat stimuli were of equal duration. The duration of these three intervals varied depending on the IOI condition used in the sequence.

### Task procedures

Participants performed two tasks, namely, a duration task and a tempo task. In the duration task, the participants were required to judge whether the comparison interval stimulus was shorter or longer than the standard interval stimulus preceding it within the same trial sequence. In the tempo task, the participants were required to judge whether the comparison beat stimulus was slower or faster than the standard beat stimulus preceding it within the same trial sequence. To enable the differentiation of the interval stimulus from the beat stimulus, the target of the tempo task was the tempo and not the duration of the beat. Nevertheless, the tempo of beats depends on the durations of IOIs of beats.

After receiving task instructions and performing practice tasks, the participants performed the tasks in the MRI scanner. The time course of a single block is shown in Figure 2. The block began with a task cue indicating whether the task was to judge duration or tempo. The task cue was Japanese words meaning length or tempo and was displayed on an LCD monitor visible via a mirror. One second after cue onset, the auditory sequence was presented via an MRI-compatible headset (SereneSound, Resonance Technology Inc). At the end of the sequence, the display switched to a response cue, requiring the participants to respond by pressing the button of a response pad within 2 s. The participants used their right index finger to indicate whether the comparison stimuli were shorter or slower than the standard stimuli, and they used their right middle finger to indicate whether the comparison stimuli were longer or faster than the standard stimuli. After the response period, a fixation cross was displayed during a 9-s intertrial interval used as a rest period.

**Fig. 2.**
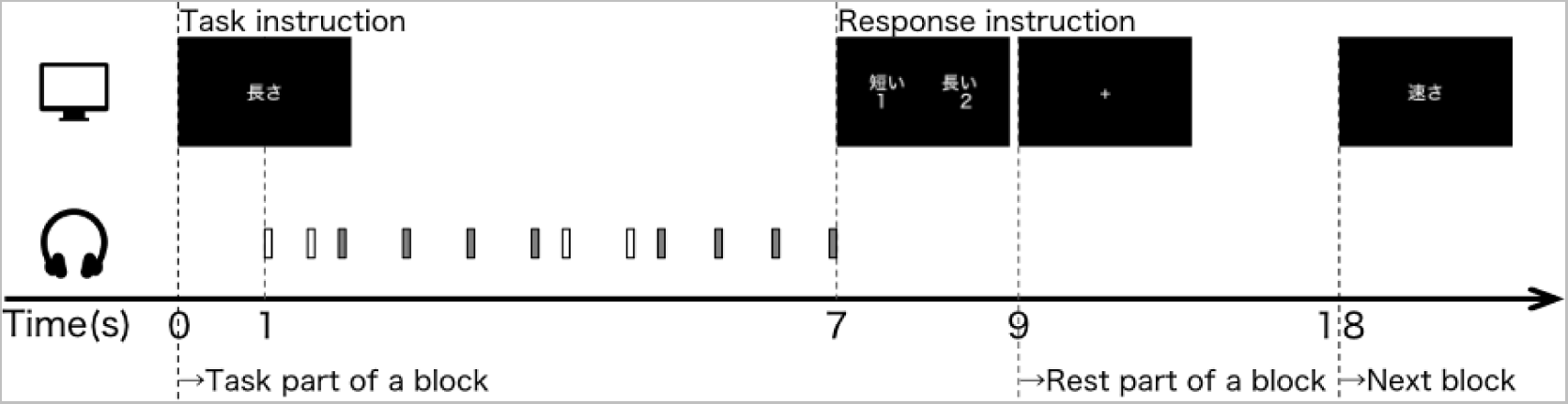
The time course of a single block. First, a Japanese word was displayed on the screen as the task instruction. One second later, the auditory sequence (6 s) was presented. The response instruction was displayed in Japanese simultaneously with the end of the auditory sequence. Two seconds later, the task ended and the display changed to the fixation cross. For the 9-s rest period of the block, the fixation cross was displayed.

The participants were instructed that the frequencies of the intervals and beats changed randomly between high and low frequencies and that the order of the intervals and beats (which stimulus was presented first) varied randomly. In addition, they were also instructed to pay attention only to the intervals during the duration task and the beats during the tempo task and to ignore the intervals during the tempo task and the beats during the duration task.

The participants underwent three sessions in total. Each session was composed of 32 blocks (16 stimuli × 2 tasks), and the order of the blocks was randomized in each session. One session lasted approximately 9.5 min. Before the first session for each participant, the sound level of the stimulus was adjusted so that the participants could comfortably and clearly hear the clicks on the background noise of the MRI scanner; the sound level was then fixed for the three sessions for that participant. A T1-weighted anatomical scan was acquired after the three sessions. The experiment was conducted consecutively in one day and took approximately 70 to 90 minutes to complete, including explanations and all scans.

### fMRI data acquisition

All fMRI scans were obtained at Hokkaido University using the same equipment, i.e., a Siemens (Erlangen, Germany) 3-Tesla Prisma scanner equipped with a 20-channel head coil. T2*-weighted echo-planar imaging (EPI) was used to acquire a total of 292 scans per session, with a gradient echo EPI sequence. The first three scans within each session were discarded to allow T1 equilibration. The following scanning parameters were used: repetition time (TR), 2000 ms; echo time (TE), 30 ms; flip angle (FA), 90 deg; field of view (FOV), 192 × 192 mm^2^; matrix, 94 × 94; 32 axial slices; and slice thickness, 3.50 mm with a 0.875-mm gap. T1-weighted anatomical imaging with an MP-RAGE sequence was performed using the following parameters: TR, 2300 ms; TE, 2.32 ms; FA, 8; FOV, 240 × 240 mm; matrix, 256 × 256; 192 axial slices; and slice thickness, 0.9 mm without a gap.

### fMRI data processing

Image preprocessing was performed using the SPM12 software (Welcome Department of Cognitive Neurology, http://www.fil.ion.ucl.ac.uk/spm). To adjust for motion artifacts, all fMRI images were subjected to an initial volume-based realignment by co-registering images using rigid-body transformation to minimize the squared differences between volumes. The realigned images were then co-registered with their T1-weighted anatomical counterparts. Finally, these images were spatially normalized using affine and nonlinear registration with the Montreal Neurological Institute (MNI) template (SPM normalization). The co-aligned, co-registered, and normalized images were resampled into 3-mm^3^ voxels (sinc interpolation) and spatially smoothed using a Gaussian kernel (6 × 6 × 6 mm^3^ full width at half-maximum).

### Statistical analysis of fMRI data

fMRI data were analyzed using the general linear model (GLM). Each trial of the duration or tempo task within a session was modeled as a separate boxcar regressor that was convolved with a canonical hemodynamic response function. Low-frequency noise was removed using a high-pass filter with a cutoff period of 128 s. Serial correlations among successive scans were estimated using an autoregressive model implemented in SPM12. To compare the regions associated with top-down processing of interval versus beat, we conducted mass-univariate analysis and created four difference images: (i) duration versus baseline, (ii) tempo versus baseline, (iii) duration versus tempo, and (iv) tempo versus duration.

Difference images of each participant were generated using a fixed-effects model and analyzed using a random-effects model with one-sample *t-*test. Activation was reported with a voxel-level threshold of *p* < 0.001 uncorrected for multiple comparisons and a cluster-level threshold of *p* < 0.05 corrected for family-wise error (FWE).

In addition, we conducted a region of interest (ROI) analysis to investigate whether top-down processing of interval in the duration task was associated with duration-based systems, and conversely, whether processing beat in the tempo task was associated with beat-based systems. The ROIs were defined as spheres of 10 mm radius around the peak voxel as reported earlier (Teki et al. 2011). We selected 10 ROIs located in the bilateral cerebellum (covering the dentate nucleus, cerebellum, vermis, cerebellar lobule IX, and cerebellar lobule X) to capture the duration-based system and 4 ROIs located in the bilateral basal ganglia (covering the caudate nucleus and putamen) to capture the beat-based system. The coordinates of each ROI are listed in Table 1. Averaged parameter estimates (beta values) for the two tasks within each ROI were obtained from the results of the fixed-effects model for each participant and compared between the two tasks using paired *t*-test.

**Table 1:**
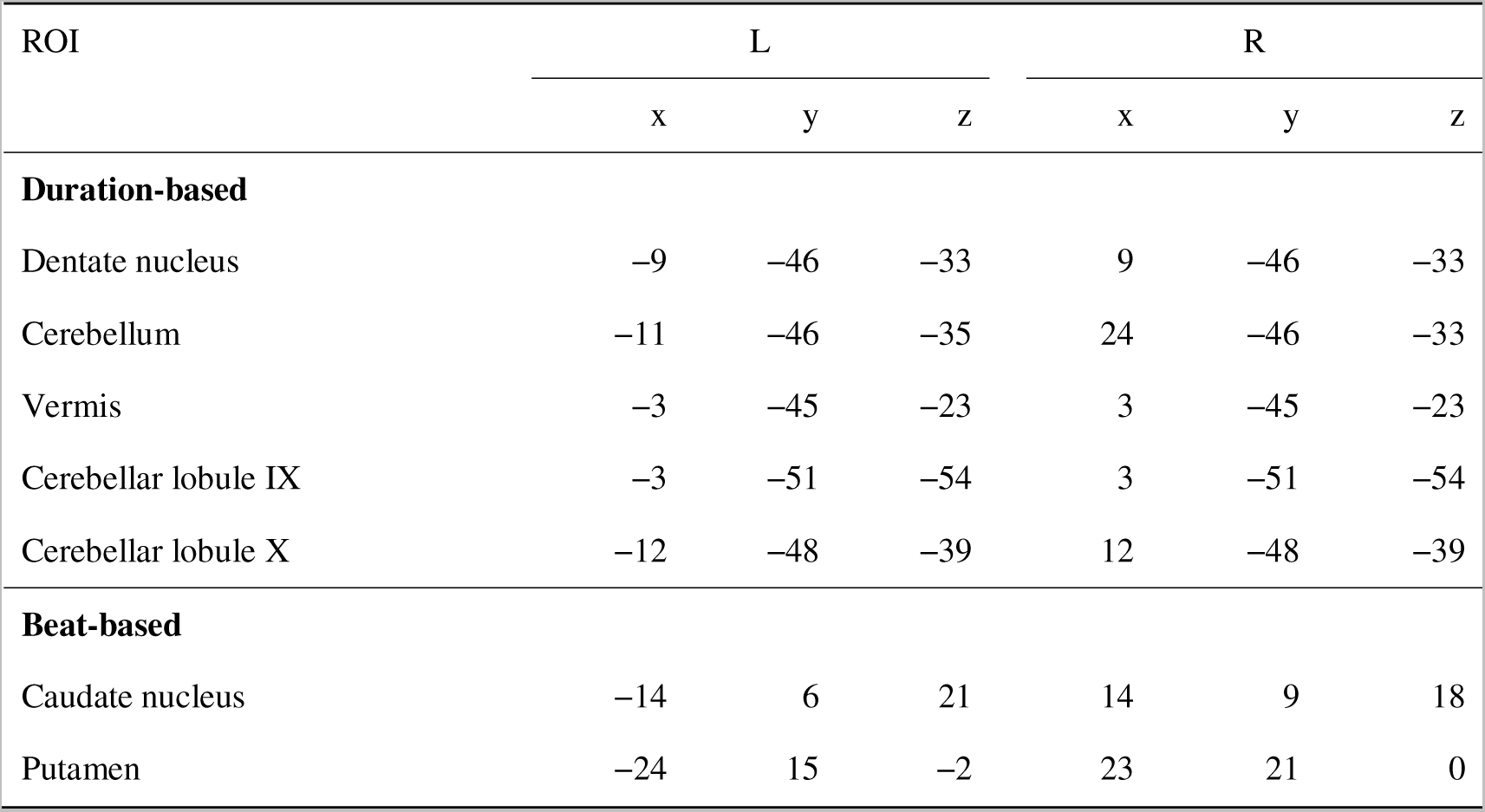
ROI coordinates.

| ROI | L |  |  | R |  |  |
| --- | --- | --- | --- | --- | --- | --- |
|  | x | y | z | x | y | z |
| **Duration-based** |  |  |  |  |  |  |
| Dentate nucleus | −9 | −46 | −33 | 9 | −46 | −33 |
| Cerebellum | −11 | −46 | −35 | 24 | −46 | −33 |
| Vermis | −3 | −45 | −23 | 3 | −45 | −23 |
| Cerebellar lobule IX | −3 | −51 | −54 | 3 | −51 | −54 |
| Cerebellar lobule X | −12 | −48 | −39 | 12 | −48 | −39 |
| **Beat-based** |  |  |  |  |  |  |
| Caudate nucleus | −14 | 6 | 21 | 14 | 9 | 18 |
| Putamen | −24 | 15 | −2 | 23 | 21 | 0 |

## Results

### Behavioral analysis

We analyzed task accuracy and reaction time separately for the duration and tempo tasks. Only the first button presses made within the 2-s response period were recorded. Two participants were excluded from behavioral analysis because of defects found in the recordings of their responses (in all sessions for one participant and in one session for the other participant). Ultimately, task accuracy and reaction time were analyzed for the remaining 18 participants. The average task accuracy was 77.66% ± 16.74% for the duration task and 77.43% ± 15.16% for the tempo task (n = 18, for both) (Fig. 3a). There was no significant difference between the two tasks (paired *t*-test, *df* = 17, *t* = 0.120, *p* = 0.906, *d* = 0.028).

**Fig. 3.**
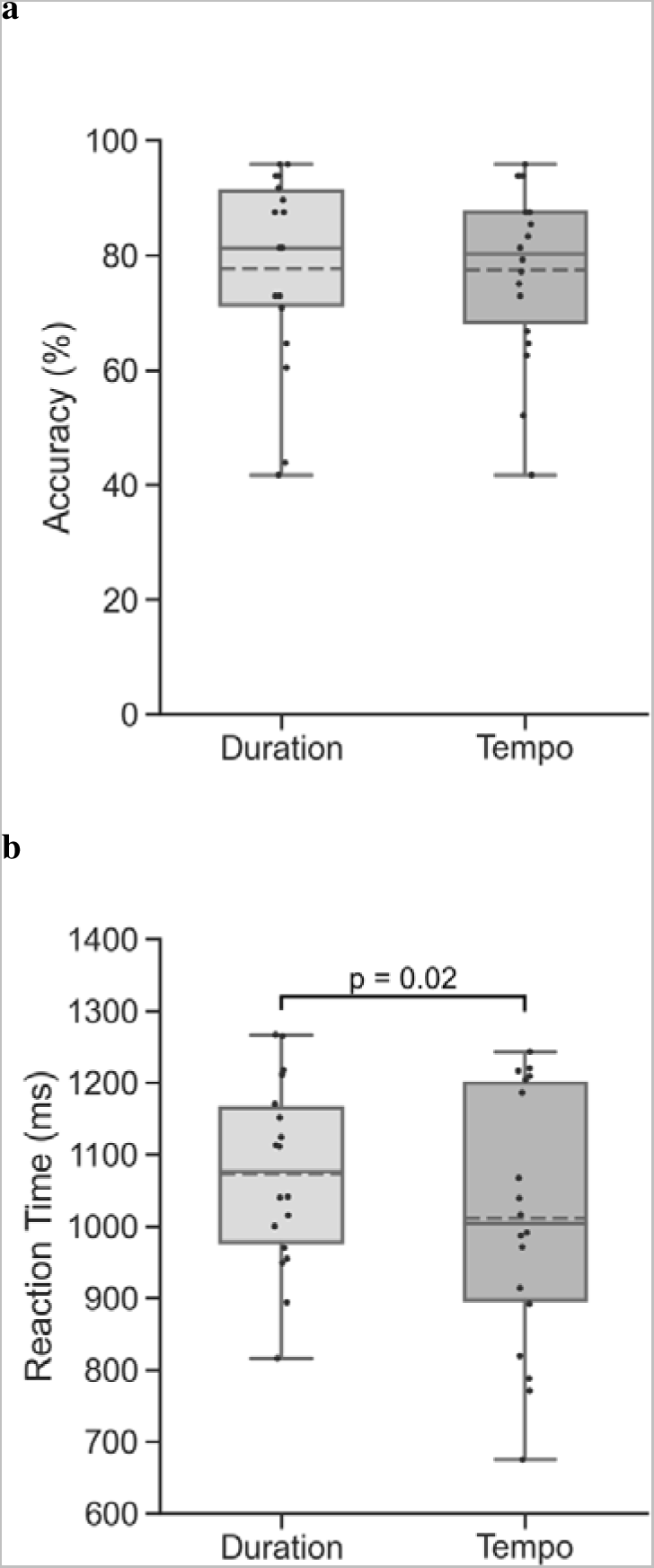
Behavioral results obtained for the duration task and tempo task. (a) Accuracy. (b) Reaction time. Each dot indicates the average result for each participant. Plots indicate the median (horizontal line inside the boxes), mean (dashed line), interquartile (25^th^–75^th^ percentile) interval (lower and upper edge of the box, respectively), and minimum and maximum values observed within 1.5 interquartile range below the 25^th^ percentile or above the 75^th^ percentile (whiskers) in the group. The significance of the difference between the tasks is indicated by p-value (n = 18).

The average reaction time was 1073 ± 129 ms for the duration task and 1012 ± 177 ms for the tempo task (n = 18, for both) (Fig. 3b). The average reaction time was significantly shorter for the tempo task than that for the duration task (paired *t*-test, *df* = 17, *t* = 2.480, *p* = 0.024, *d* = 0.585).

### fMRI data analysis

First, we directly compared the brain regions associated with the duration task and tempo task to detect possible differences in brain networks processing intervals and beats. However, two difference images, i.e., duration versus tempo and tempo versus duration, indicated that there was no significant difference between the activated regions during the two tasks (n = 20). We then repeated the same analysis but only included participants who performed the tasks with above-chance accuracy. Seventeen participants whose task accuracy was over chance (50%) were selected using a binomial test (*p*s < .01). The task accuracy was measured for both tasks as a whole and was calculated as an average across all 96 trials of the three sessions for each participant, except for the two participants who were excluded from the behavioral analysis because of defects in the recordings of their responses. The accuracy for one participant was calculated based on the results of two accurately recorded sessions, and the other participant was excluded from this analysis. Nevertheless, the two difference images, i.e., duration versus tempo and tempo versus duration, of the 17 participants indicated no significant difference in the activated regions during the two tasks. In brief, we found no evidence that interval processing and beat processing activated different regions with our tasks.

Next, we separately compared the brain activity during the duration or tempo task to the brain activity at the baseline to identify the brain regions associated with either task (Tables 2 and 3). The difference images, i.e., duration versus baseline and tempo versus baseline, for the above 17 participants revealed that similar regions were activated during the two tasks (Fig. 4). The regions that were activated during both the duration task and tempo task, shown in blue in Fig. 4, included the cerebellum and basal ganglia, which are essential components of the proposed duration-based and beat-based systems, respectively. These results indicated that the brain regions activated by the two tasks overlapped to a great extent, with the overlap including the cerebellum and basal ganglia.

**Fig. 4.**
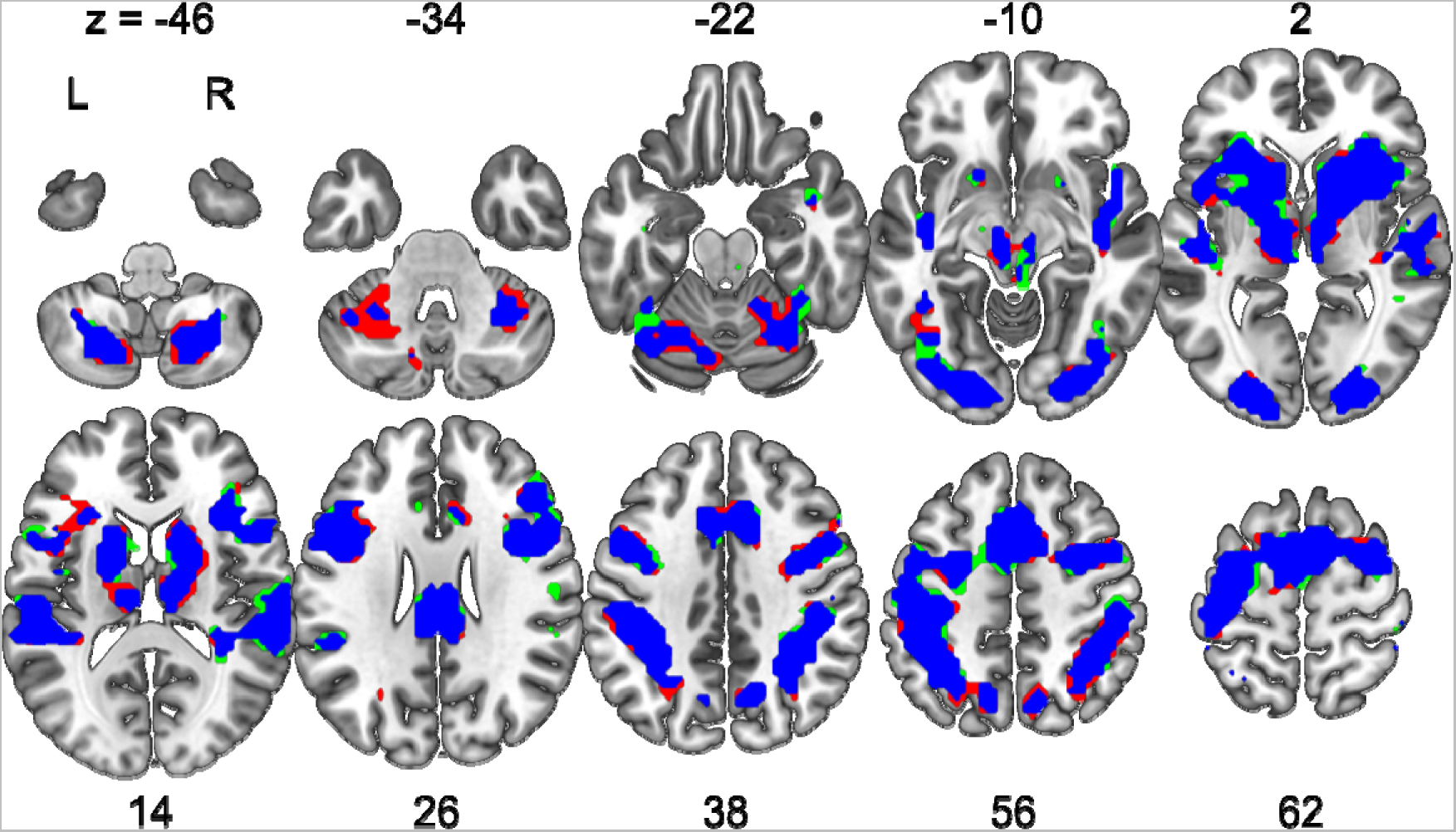
Activation of brain regions observed in fMRI scans during the tasks (levels relative to baseline). Activation by the duration task (red), tempo task (green), or both tasks with spatial overlap (blue) is reported with a voxel-level threshold of *p* < 0.001 uncorrected for multiple comparisons and a cluster-level threshold of *p* < 0.05 corrected for family-wise error (FWE). L, left hemisphere; R, right hemisphere. The MNI Z-coordinate is indicated for each successive horizontal cut. The MNI coordinates of the activated foci are reported in Tables 2 and 3.

**Table 2:**
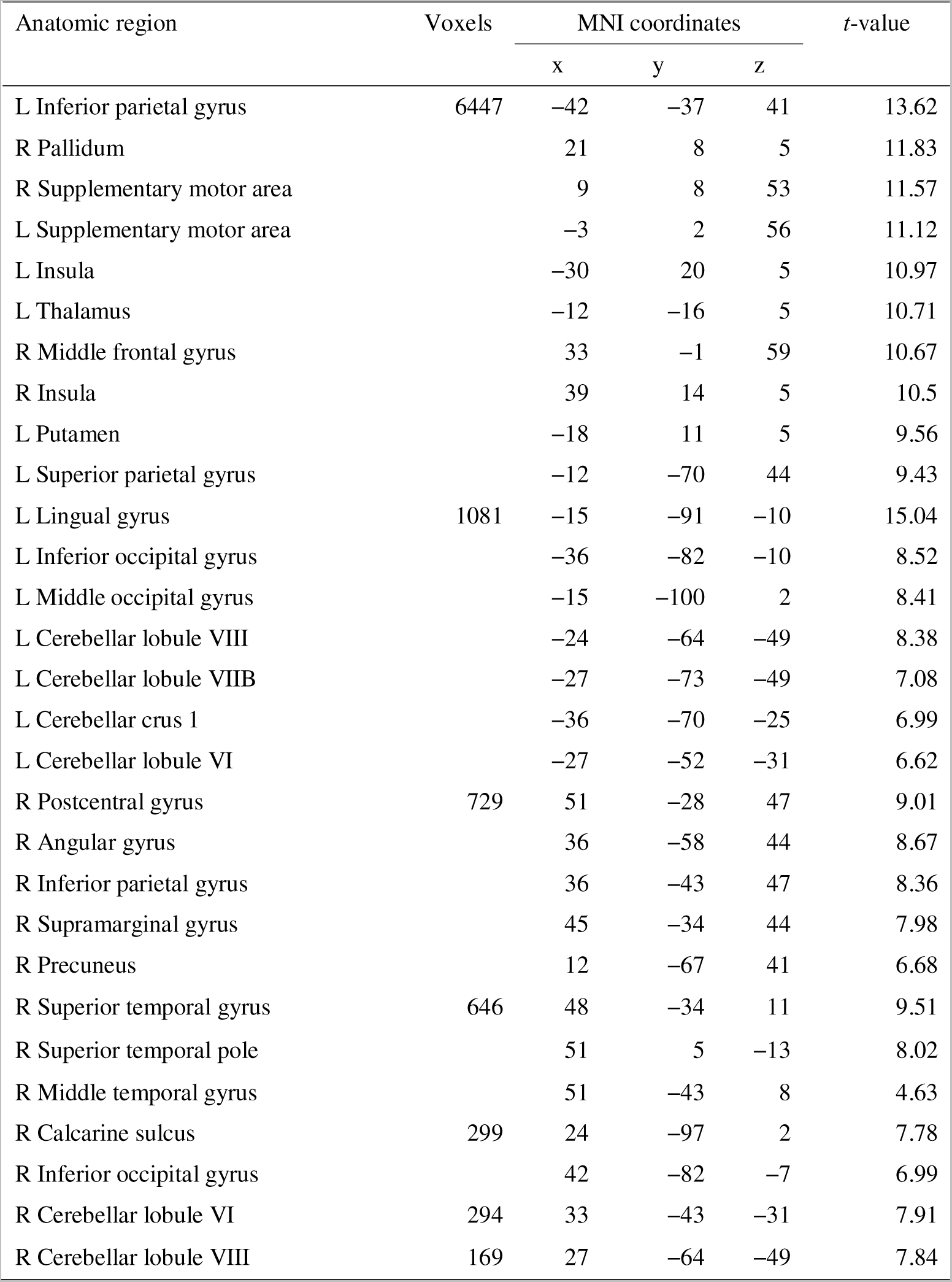
Brain activation observed in the duration task compared with the baseline. List of the anatomical regions, peak voxel coordinates, and *t*-values for each activated area.

**Table 3:**
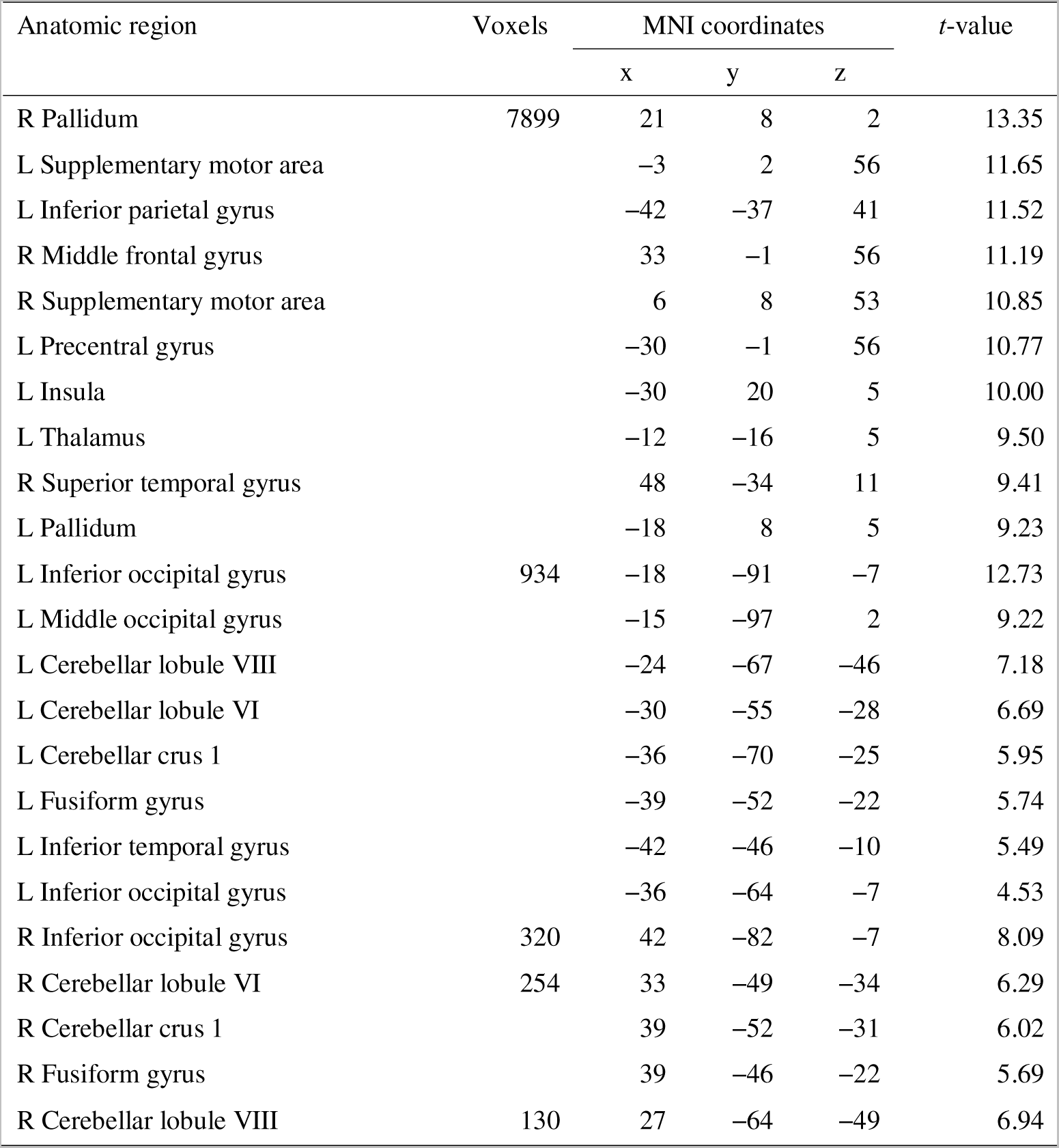
Brain activation observed in the tempo task compared with the baseline. List of the anatomical regions, peak voxel coordinates, and *t*-values for each activated area.

Finally, we compared the parameter estimates (beta values) for the two tasks within each ROI in the cerebellum and basal ganglia to determine whether the duration- and beat-based systems were involved in top-down processing of interval and beat. The average beta values in each ROI of the cerebellum showed no significant difference between the two tasks (*t*-test, *df* =16, *ps* > 0.15) (Fig. 5a). A similar analysis within ROIs covering the basal ganglia yielded the same results (*t*-test, *df* =16, *ps* > 0.13) (Fig. 5b). Thus, with this analysis too, we found no difference in the brain activations during the two tasks, especially in the cerebellum and basal ganglia.

**Fig. 5.**
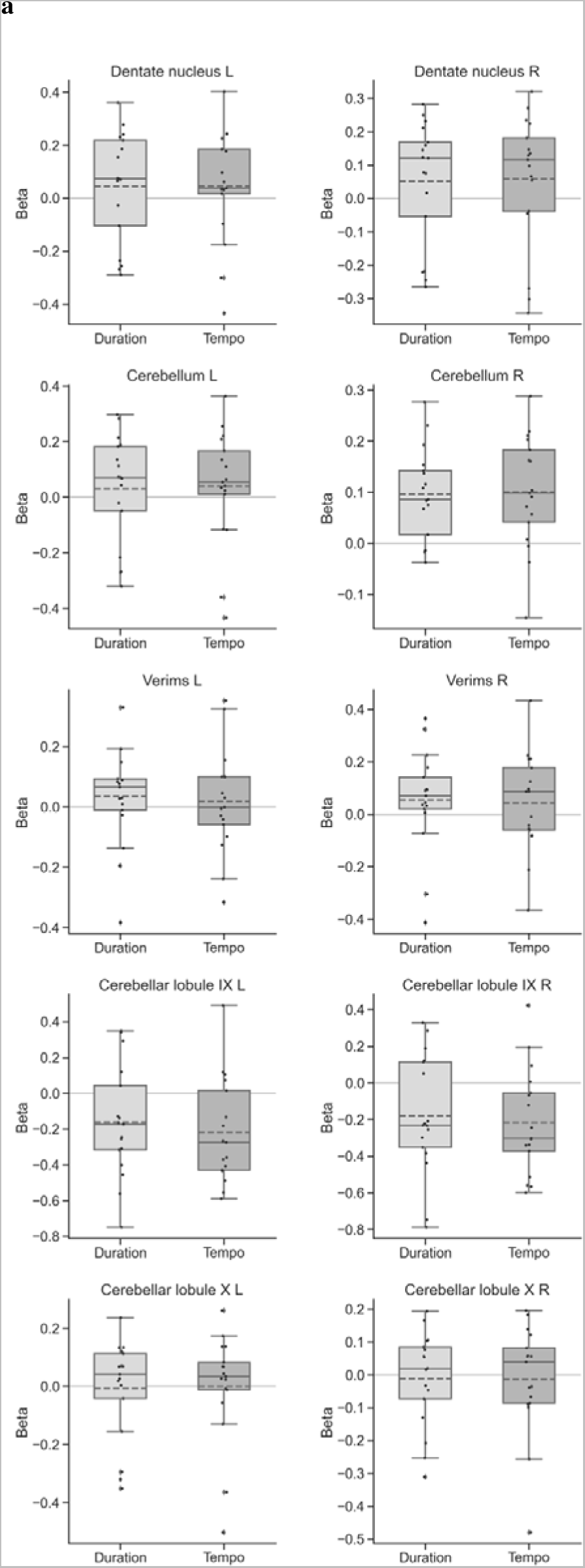
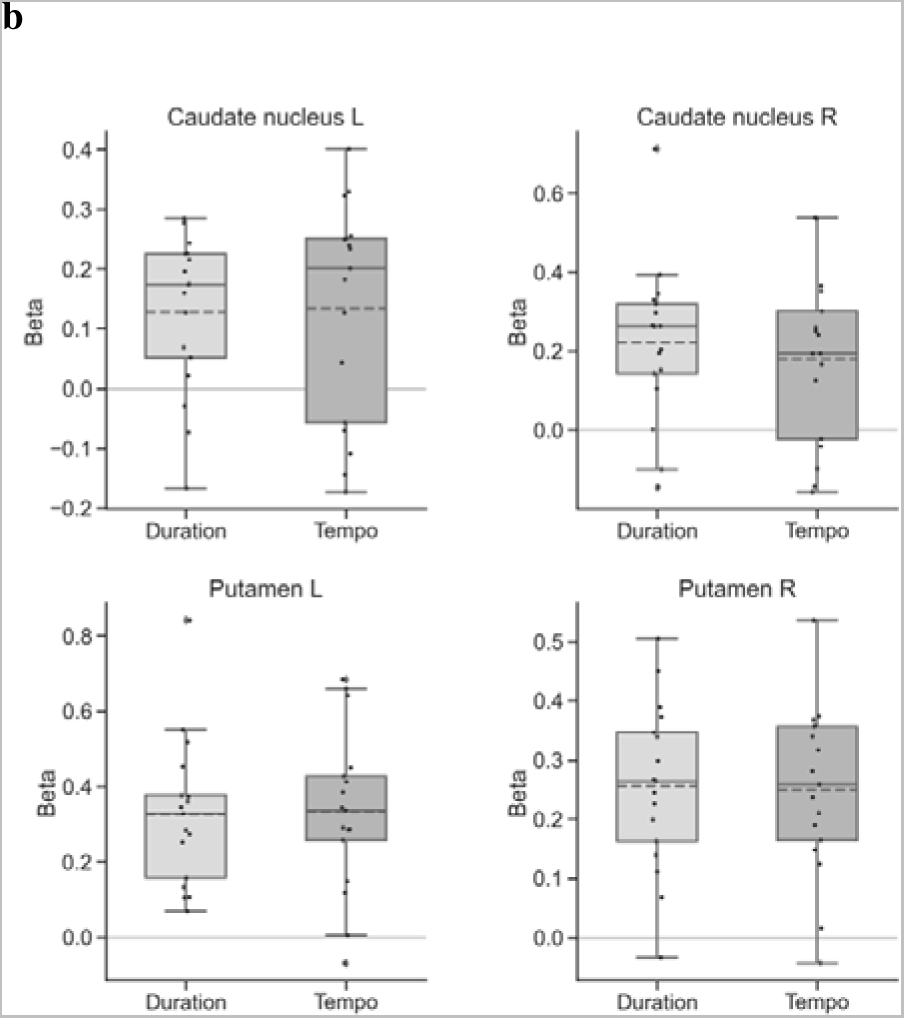
Averaged parameter estimates (beta values) for the duration task and tempo task. Estimates were based on activations in the (a) ROIs in the cerebellum and (b) ROIs in the basal ganglia. L, left hemisphere; R, right hemisphere. Each dot indicates the average result for each participant. Plots indicate the median (horizontal line inside the boxes), mean (dashed line), interquartile (25^th^–75^th^ percentile) interval (lower and upper edge of the box, respectively), and minimum and maximum values observed within 1.5 interquartile range below the 25^th^ percentile or above the 75^th^ percentile (whiskers) in the group. Outliers are marked by the rhombus symbol.

## Discussion

The objectives of this study were to (a) compare brain activation during duration-based timing versus beat-based timing under top-down processing and (b) examine how top-down processing of these two timings affected activations in the olivocerebellar network and striato–thalamo–cortical network. To address these questions, we conducted an fMRI experiment in which participants were asked to distinguish the duration of single intervals or the tempo of beats. The sets of auditory sequences used in the two tasks were identical by design to equalize the bottom-up inputs while processing the duration-based and beat-based timings. In addition, the task instructions were designed to elicit different top-down processing, such as attention and strategy, for the duration-based and beat-based timings; the participants were instructed to attend only to the single intervals during the duration task and to the beats during the tempo task.

To compare brain activation involved in top-down processing of the two timing tasks, it is necessary to balance task difficulty as it affects neural activation (Gould et al. 2003; Zhang et al. 2018; Brechmann and Angenstein 2019), even in timing tasks (Tregellas et al. 2006; Livesey et al. 2007; Lewandowska et al. 2010). Using two tasks that were balanced for difficulty in our experiment, we found no significant difference between the average task accuracies for the duration versus tempo tasks.

In contrast, the reaction times were significantly different between the duration and tempo tasks. The speed–accuracy tradeoff predicts that reaction times of choice responses covary with their accuracy (Wickelgren 1977; Heitz 2014). This result was inconsistent with our results—while reaction times of the two tasks were significantly different, the average accuracies were equivalent. Our results suggest that although the stimuli presented in the two tasks were identical, the participants used different strategies to respond. In other words, we were likely comparing two different behaviors in response to identical sets of stimuli used in the duration and tempo tasks, implying that the brain activation levels measured during the two tasks presumably reflected top-down processing of both duration-based and beat-based timings.

The mass-univariate analysis revealed that the task-contrasting difference images, i.e., duration versus tempo and tempo versus duration, showed no significant difference in brain activation between the duration task and beat task. ROI analysis of the parameter estimates in the basal ganglia and cerebellum also revealed no difference between the two tasks. These results suggest that the same brain regions were activated during interval processing and beat processing when identical auditory stimuli were used in the two tasks. However, in this experiment, performing the two tasks required paying attention to either the intervals or the beats in the sequences, and the behavioral results suggested that the participants’ response behavior was also different between the two tasks. Our results were not consistent with those of previous studies that revealed the involvement of distinct neural substrates in perceptual processing of duration-based and beat-based timings. However, those experiments used auditory stimuli with different properties for each kind of timing task (Grube et al. 2010b, a; Teki et al. 2011). It is possible that the distinct neural systems identified as being involved in perceptual processing of duration-based timing and beat-based timing were stimulus-driven and not affected by top-down processing, such as attention and strategy elicited by the task instruction. In comparison, our results suggest that at least in terms of brain activation levels, top-down processing of the two types of perceptual timing systems may involve a common neural basis.

Our results support the neural model of time perception first proposed by Teki, Grube, and Griffiths (2012). In this model, the beat-based striato–thalamo–cortical network and the duration-based olivocerebellar network are integrated as a single system. The cerebellum and basal ganglia are connected to multiple areas of the cerebral cortex through multisynaptic loops and may work in parallel to time a wide range of durations (Meck 2005; Allman and Meck 2012). Moreover, the cerebellum is interactively connected to the basal ganglia via disynaptic pathways, and this connection may make the cerebellum and basal ganglia an integrated functional network (Hoshi et al. 2005; Bostan et al. 2010; Bostan and Strick 2010). The model proposes that the olivocerebellar circuits complement the timing measurements performed by the striato–thalamo–cortical circuits by performing an error correction, and that each of these circuits is activated depending on the temporal predictability of the stimulus, such as isochronism versus irregularity. In our experiment, the stimuli of both the duration and tempo tasks were the same sequences, and the sequence contained both isochronous beats and single intervals. Although the participants were instructed to pay attention only to the target part and to ignore the non-target part, our data suggest that the whole sequence, including the part not required to perform the tasks, was processed collectively.

In our experiment, the standard intervals were fixed at 400 or 600 ms. The participants could have memorized each of these durations during each session and compared this internalized reference to the comparison stimulus, potentially eroding the application of the different strategies for the two tasks. Indeed, Miller and McAuley (2005) and McAuley and Miller (2007) showed that tempi of standard stimuli are memorized and that the internalized reference affects timing accuracy by the end of blocks; however, the internalized reference is formed by averaging each standard tempo throughout an experiment, rather than by memorizing individual standard tempo. The authors demonstrated that time estimation is more accurate when the tempo of a standard stimulus matches the average tempo of standard stimuli across the IOI condition, rather than a specific tempo. The IOI condition that we used was two conditions, and the average duration of these IOIs did not match either of these IOI conditions. Thus, although the internalized reference could still have influenced the time estimations, the participants likely processed the actual IOIs presented on each trial, and the strategies for the two tasks may not be equalized by the use of the internalized reference.

The participants may have discriminated the tempo of the beat stimuli based on only one or two intervals and prepared their responses before the end of the stimulus presentation. This possibility could explain the difference in reaction time between the duration task and tempo task. Nevertheless, it is suggested that three intervals are required for tempo comparison. Pfeuty et al. (2003) recorded electroencephalogram (EEG) and compared the contingent negative variation (CNV) during tempo encoding and comparison using three- or six-interval isochronous stimuli. CNV amplitude increased up to the third interval during tempo encoding in both the three- and six-interval conditions and was sustained during tempo comparison in the three-interval condition. The authors suggest that three intervals are a critical limit for building a memory trace of successive intervals and that the subjects encoded a new memory trace during the tempo comparison of three intervals. Accordingly, our participants may have attended to all three intervals and not prepared their responses immediately after the first or second interval.

In addition, the participants may have applied the same strategy to both the duration and tempo tasks, by comparing durations of single intervals in the standard and comparison beats during the tempo task. This strategy could provide the participants with time to prepare their responses during the remaining two intervals for the tempo task. The possibility that the participants used this same strategy to focus only on durations for both tasks could explain the difference in response time and the absence of significant differences in brain activation between the two tasks. However, the task procedures probably prevented the participants from using this strategy. The participants were required to indicate the shorter comparison interval by pressing the left button with their right index finger for the duration task, and the faster comparison beat (i.e., equivalent to a shorter duration) by pressing the right button with their right middle finger for the tempo task. In other words, the participants had to use the opposite button for the two tasks to indicate the shorter durations. Given that the two tasks were presented in random order during each session, participants who used the duration-only strategy must have had to convert the button position for the tempo task. This additional conversion would have reduced the time available for participants to prepare their responses before the end of the stimuli presentation. Furthermore, the difference in the IOI ratio between the two tasks may also have made it difficult for the participants who used the duration-only strategy to prepare their responses before the end of the stimuli presentation. Because the IOI ratio of the comparison stimuli was smaller for the beat stimuli than for the interval stimuli, discriminating the duration of the beat stimuli was more difficult than discriminating the duration of the interval stimuli. The participants could have used the second and third intervals of the beats to improve their accuracy in discriminating the duration of the beat stimuli. However, this would have shortened the time to prepare the responses before the end of the stimuli presentation. To sum up, considering the button position conversion and the difference in the IOI ratio, it is likely that the participants did not use the duration-only strategy and used the different strategy for the two tasks, according to the task instructions.

One limitation of this study was the imbalance of the stimuli within the sequences. Specifically, the single intervals and isochronous beats were arranged in series in an alternating order, and the standard stimuli were separated from the comparison stimuli in time. These sequence designs were aimed at making sequences containing both single intervals and isochronous beats and balance them for the two tasks to make the participants keep listening to the whole sequence. However, this sequence design did not balance the intervals and beats. For example, the proportions of lengths within the sequences are different between the intervals and beats. Furthermore, because of the series arrangement, the participants might have also paid attention to the beats as much as to the intervals during the duration tasks, and vice versa. Future experiments need to avoid these problems by developing stimuli and tasks in which regular and irregular sequences are presented and processed in parallel.

In summary, we found that neural activations during discrimination of single intervals and isochronous beats were not different when the stimuli contained both intervals and beats. Our results suggest that top-down processing of perceptual duration-based timing and perceptual beat-based timing have a common neural substrate. Our research provided new insights into the distinct and interconnected nature of duration-based and beat-based timing systems, which have been studied using stimuli with different regularities by separating bottom-up and top-down processing through the equalization of auditory input and the differentiation of task instruction.

## Statements and Declarations

### Ethical standards

The experimental protocol received approval from the local ethics committee, and the experiments were conducted in accordance with the 1964 Declaration of Helsinki. Written informed consent was obtained from each participant.

### Author contributions

Mitsuki Niida: Conceptualization, Methodology, Software, Formal analysis, Investigation, Data Curation, Writing - Original Draft, Visualization. Yusuke Haruki: Investigation, Data Curation, Writing - Review & Editing. Fumihito Imai: Investigation, Data Curation, Writing - Review & Editing. Kenji Ogawa: Conceptualization, Methodology, Software, Formal analysis, Resources, Writing - Review & Editing, Supervision, Project administration, Funding acquisition.

### Data availability

The datasets generated during and/or analyzed during the current study are available from the corresponding author on reasonable request.

## Acknowledgments

The authors would like to thank Enago (www.enago.jp) for the English language review.

